# Benchmarking honorific academic recognitions against citation metrics

**DOI:** 10.64898/2026.11.11.766046

**Authors:** John P.A. Ioannidis, Jeroen Baas

## Abstract

We examined how diverse metrics track individuals receiving major honorific recognitions and whether higher, more prestigious recognitions are associated with higher coverage among a database of top-cited scientists. We assessed with Scopus data the newly inducted (in 2026) 2,038 members of 14 major academies and honorific societies, endowed professors at Stanford University (n=773) and Michigan State University (MSU) endowed professors (n=129) and distinguished professors (n=112), and Nobel (n=166) and Turing (n=25) laureates awarded in 2010-2026 (n=191) for total publications, total citations, h-index, coauthorship-adjusted hm-index, citations to single-, single/first-, and single-/first-/last-authored (SFL) publications, and a composite citation indicator (c-score). For those with ≥5 full papers and ≥20 published items indexed in Scopus, metrics were mapped as percentile values among authors sharing the same primary scientific subfield and ≥5 full papers. Citations to SFL publications had most frequently the best median percentile values and c-score had the best median of median percentile values (0.81%). Subfield-adjusted median percentile values were 0.04-0.23% for Turing/Nobel, 0.23-3.01% for academies and honorific societies, 2.74% for MSU endowed professors, 1.29% for MSU distinguished professors, 0.94% for Stanford endowed professors for citations to SFL publications; and 0.04-0.26%, 0.36-3.85%, 3.21%, 1.55%, 1.03%, respectively, for c-score. The percentages of honorees included in the Stanford/Elsevier listing of top-cited scientists (top-2% of c-score in their scientific subfield or top-100,000 in c-score across all science) followed the perceived prestige gradient: 99% for Turing/Nobel laureates, 84-92% for the national US academies and the UK Royal Society as well as AAP and ACM, 51-76% for other academies/societies, 78% for Stanford endowed professorships, 72% for distinguished MSU professorships, 52% for MSU endowed professorships. Several honored scientists had at least one retraction (highest for medicine (17.5%) and chemistry (11.4%) Nobel laureates), and few had very high proportion of papers in Scopus-discontinued titles, while high self-citations were rarely encountered. Metrics may influence honors and vice versa. The current evidence suggests strong tracking of the median values, especially for citations to SFL publications and c-score, against major recognitions; and a gradient of coverage in a list of top-cited scientists congruent with the perceived relative prestige of these recognitions.

---

Citation metrics are widely used and misused to assess scientific impact (1). Some of the most popular metrics, such as the h-index, were originally proposed and promoted with a validation that they can have very high values among scientists who have received major honorific recognitions, e.g. Nobel prizes and membership in the US National Academy of Sciences (2). The ability to specifically predict extreme honors such as Nobel prizes has been debated but probably remains limited (3–5). The careers of these exceptional people are multifaceted (6) and thus not easy to put into single numerical metrics. Moreover, a very large number of scientists may have performed exceptional work, but only 1 to 3 get the most prestigious awards such as Nobel, Turing, Fields, or Abel. However, for other major recognitions there may be far more slots for honorees available. For example, several major academies and honorific societies elect many dozens or even hundreds of new members every year; and major universities may have hundreds of endowed professorships. These honors are still highly prestigious and selective, but not as rare. It would be interesting to assess how a diverse set of citation metrics perform in benchmarking the cohorts of scientists who receive these honorific recognitions. Do these metrics track the relative perceived prestige of these recognitions? Are many, most or all scientists who are given such honors top-cited authors? Which metrics have the best values for the average person who is offered different coveted honors when placed against peers in the same scientific discipline?

We aimed to address these questions using data from recently inducted members from 32 groups: 14 prestigious academies and honorific societies, Nobel and Turing laureates and endowed professors in various disciplines from two major universities. We explored diverse citation metrics that focus on overall productivity, overall citation impact, co-authorship-adjusted impact, citation impact for publications in specific influential authorship positions, and a composite citation indicator (7). We also noted whether these prestigiously recognized individuals had any evidence of proxies that may highlight potential issues with research integrity (8) in their records.

## METHODS

### Evaluated cohorts of scientists with honorific recognitions

We considered the newly inducted (in 2026) members of 32 groups of honorees: 14 prestigious academies and honorific societies, science-related Nobel and Turing laureates, and the endowed professors in diverse disciplines of Stanford University (a top, private university ranked globally #2 by QS ranking (9) and #3 by overall CWUR ranking (#3 in Research Rank) (10)) and of Michigan State University (MSU, a very good large public university ranked globally #182 by QS ranking (9) and #168 by overall CWUR ranking (#131 in Research Rank) (10)).

We considered academies and honorific societies that have a selective induction process where no self-nomination is allowed; include at least 10 living Nobel or Turing laureates as regular members (as documented by searching their websites or online searches); and inducted more than 60 new members in 2026: US National Academy of Sciences (NAS), US National Academy of Engineering (NAE), US National Academy of Medicine (NAM), American Academy of Arts and Sciences (AmAAS), Academia Europaea (AE), European Academy of Sciences and Arts (EASA, 2025-2026 fellows, announced as one list in 2026), American Association for the Advancement of Science (AAAS, 2025 fellow class announced in 2026), American Academy of Microbiology (AAM), Association of American Physicians (AAP), American Society for Clinical Investigation (ASCI), Association for Computing Machinery (ACM, 2025 fellow class announced in 2026), European Molecular Biology Organization (EMBO), UK Royal Society (RS), and Royal Society of Canada (RSC). For Turing awards and Nobel prizes in Chemistry, Physics, Medicine, and Economics laureates, we included those recognized and given the awards in the years 2010-2026 (Turing for 2026 will be announced in 2027). For Stanford, we considered all endowed professorships as of October 2026 (excluding endowed junior positions, scholars, and directors) and used data separately for each School with at least 20 such professorships (Business, Education, Engineering, Humanities and Sciences, Law, Medicine, Sustainability). Similarly, for MSU we also used data as of October 2026 separately for each College with at least 20 such professorships (Business, Engineering, Natural Science) and we also considered an additional set, the MSU Foundation University Distinguished and John A. Hannah Distinguished Professors. Supplementary Table 1a identifies the publicly available online lists, and Supplementary Table 1b shows the names as they appeared in those lists.

For some of these honors, the recognized impact may not necessarily focus primarily on work reflected by scholarly publications. E.g. academies and honorific societies may occasionally elect members primarily or exclusively for their excellence in entrepreneurship, social service, educational impact, or other types of merit, e.g. some academies also include artists. We therefore included in the metrics analysis only those members in each cohort who had at least 5 Scopus-indexed publications that were full original articles, reviews, or conference papers (“full papers”) and at least 20 total Scopus-indexed published items of any type.

### Crosslinking honored scientists to Scopus author ID files

The list of honorary awardees was compiled from source materials (Supplementary Table 1a) and linked to Scopus Author IDs in a multi-step process. Each entry was first assessed by Claude Haiku 5.5 to determine whether it referred to a person and to extract usable information, including the name, title, department, institution, location, and research area (Supplementary Table 1b). Institution names and variants were used to identify candidate Scopus affiliation IDs. We then searched Scopus for author candidates using a series of queries that combined name information with affiliation IDs, institution, and location, followed by a broader name search.

Candidates were ranked using evidence from name, affiliations, institution, location, subject area, and publication years. Claude Haiku 5.5 assessed the leading candidates for inclusion, exclusion, or further review. If there were no high-confidence results, or the decided “include” recommendation included a name conflict, cases are rejudged by Claude Sonnet 5.5. It was possible to link more than one Author ID, when the evidence indicated that Scopus had split an awardee’s publications across multiple profiles.

We manually reviewed the results and documented any interventions for additional corrections (Supplementary Table 2). For example, in the 2026 Nobel Chemistry reporting run, we changed the source spelling *Aleksey Yekimov* to *Aleksey Ekimov* and retained *Aleksey Yekimov* as a name variant.

### Scopus database metrics and percentile placement

We used the August 1, 2026 data freeze that has been used to generate the latest iteration of the Stanford/Elsevier database of top-cited scientists (11), an annually updated list based on Scopus data (12). We captured Scopus publications and citations up to end of 2025 for all 11,619,713 Scopus author ID files that included at least 5 full papers. For each of these authors, we generated information on their publication age (number of years since the first published item indexed in Scopus), and diverse citation metrics based on citations received in 1996-2025 (total citations, Hirsch h-index (2), Schreiber hm-index (13), citations to single-authored items, citations to single-or fist-authored items, citations to single-, first-, or last-authored items, and a composite citation indicator (c-score) that combines the previous 6 metrics, as previously described (7,14,15). Citations received before 1996 are elliptically covered in Scopus, and some of the older scientists may have had some missed citations, although it is likely that the proportion of pre-1996 citations would be minimal for the examined individuals. Citation metrics included self-citations.

Each evaluated eligible scientist was placed in a percentile for each of the examined metrics with two decimal points accuracy, considering all scientists and separately considering only the scientists classified into the same primary scientific subfield according to the Science Metrix classification (16) that divides science into 174 subfields. E.g. a value of 0.12% for a scientist’s c-score in her subfield means that only 0.12% of the scientists in the same subfield who have at least 5 full papers have better/higher c-score values. It is expected that for honorific groups where scientists work in fields with high citation densities, their percentile ranking will spuriously seem nicer when all scientists are considered rather than when only scientists in their subfield are considered. Conversely, it is expected that for honorific groups where scientists work in fields with low citation densities, their percentile ranking will spuriously seem less attractive when all scientists are considered than when only scientists in their subfield are considered. Unless stated otherwise, in the Results we focus on percentile rankings within the subfield which are more appropriately matching each scientist against their subfield peers.

### Percentage of honored scientists who are listed as top-cited

Scientists at the top-2% of the composite score in each of 174 scientific subfields and those among the top-100,000 of the composite score (either with-or without self-citations) across all science are included in the annually updated Stanford/Elsevier list of top-cited scientists (11). Top-cited scientists are catalogued separately for career-long impact and for impact in the most recent single year. For each group of honored scientists, we present the percentage that are among the latest edition (August 2026) of the top-cited scientists’ data. This is limited to members who meet the minimum productivity criteria mentioned above. We examined whether coverage percentages in the Stanford/Elsevier listing for the examined groups were higher for more prestigious groups, i.e. highest of all in the Turing and Nobel laureates, very high in the official national academies of the US and UK than in other academies and honorific societies, higher for AAP than for ASCI (brother organizations, where ASCI is the first step for relatively younger scientists and AAP for more senior ones), higher for Stanford than for MSU professorships (based on their institutional rankings), and higher for MSU distinguished professorships than for MSU professorships.

### Research integrity proxies

It is also increasingly recognized that it is important to present not only positive impact but also any evidence or markers of potential problems with research integrity (8,17,18). Therefore, we have also collected data on three research integrity proxies: number of publications that have been retracted according to Retraction Watch database (19) for reasons other than journal or publisher errors (20,21); proportion of self-citations (from all authors) among total citations (22); and number of publication in journals that lost their indexing status upon review by the Scopus review board, as in previous work on these proxies (8). Therefore, we report for each group of honored scientists whether any of them had any such retracted publications; were at the top-5% of self-citation rate for top-cited scientists in their subfield; or were at the top-5% of proportion of publications in discontinued journal titles among top-cited scientists in their subfield.

## RESULTS

### Assessed honored scientists, productivity criteria, and publication age

As shown in Table 1, the groups of newly inducted members across the 14 academies and honorific societies included a total of 2,038 individuals, of which 1,812 (88.9%) passed the productivity threshold for further analysis. 100% of newly inducted NAS members and 98.6% of EMBO members passed the productivity threshold, while the lowest percentages passing the productivity criteria were seen for AmAAS (69.4%), EASA (71.2%) and NAE (75.3%), reflecting mostly the induction of many new members for diverse types of merit other than what is reflected in scholarly publications and their impact.

**Table 1.** Groups of honored individuals, numbers eligible for analysis, and publication age.

| Honorific group | Total members | Eligible for analysis | Median age | 25th percentile | 75th percentile |
|---|---|---|---|---|---|
| American Association for the Advancement of Science | 449 | 424 | 32 | 26 | 38 |
| American Academy of Microbiology | 63 | 63 | 29 | 24 | 34 |
| Association of American Physicians | 72 | 72 | 31 | 26 | 38 |
| Academia Europaea | 254 | 227 | 31 | 24 | 38 |
| ACM | 71 | 69 | 27 | 24 | 33 |
| American Academy of Arts and Sciences | 252 | 175 | 35 | 27 | 43 |
| American Society for Clinical Investigation | 100 | 100 | 22 | 18 | 24 |
| European Academy of Sciences and Arts | 104 | 74 | 30 | 23 | 35 |
| EMBO | 71 | 70 | 25 | 20 | 31 |
| National Academy of Engineering | 158 | 119 | 39 | 33 | 44 |
| National Academy of Medicine | 100 | 99 | 32 | 26 | 38 |
| National Academy of Sciences | 145 | 145 | 38 | 31 | 45 |
| Royal Society | 94 | 90 | 36 | 31 | 40 |
| Royal Society of Canada | 105 | 85 | 30 | 26 | 35 |
| MSU endowed chairs and professors, all colleges | 129 | 108 | 28 | 22 | 37 |
| MSU endowed chairs and professors, Eli Broad College of Business | 21 | 12 | 28 | 25 | 30 |
| MSU endowed chairs and professors, College of Engineering | 20 | 19 | 34 | 23 | 37 |
| MSU endowed chairs and professors, College of Natural Science | 27 | 26 | 29 | 19 | 38 |
| MSU Foundation, University Distinguished and John A. Hannah Distinguished Professors | 112 | 105 | 35 | 27 | 43 |
| Stanford endowed professorships, all units | 773 | 672 | 34 | 28 | 42 |
| Stanford endowed professorships, Graduate School of Business | 74 | 58 | 28 | 21 | 34 |
| Stanford endowed professorships, Graduate School of Education | 23 | 23 | 32 | 23 | 34 |
| Stanford endowed professorships, School of Engineering | 84 | 79 | 36 | 30 | 43 |
| Stanford endowed professorships, School of Humanities and Sciences | 240 | 183 | 35 | 28 | 45 |
| Stanford endowed professorships, Stanford Law School | 44 | 26 | 30 | 23 | 36 |
| Stanford endowed professorships, Stanford Medicine | 245 | 243 | 34 | 28 | 42 |
| Stanford endowed professorships, Stanford Doerr School of Sustainability | 23 | 23 | 40 | 32 | 44 |
| ACM A.M. Turing Award laureates, award years 2010-2025 | 25 | 24 | 50 (43*) | 46 (38*) | 55 (47*) |
| Nobel Prize in Chemistry laureates, 2010-2026 | 45 | 44 | 55 (46*) | 45 (39*) | 59 (52*) |
| Sveriges Riksbank Prize in Economic Sciences laureates, 2010-2026 | 37 | 37 | 48 (40*) | 41 (34*) | 53 (46*) |
| Nobel Prize in Physiology or Medicine laureates, 2010-2026 | 40 | 40 | 51 (44*) | 44 (38*) | 56 (47*) |
| Nobel Prize in Physics laureates, 2010-2026 | 44 | 43 | 55 (48*) | 44 (36*) | 61 (57*) |

ACM: Association for Computing Machinery; EMBO: European Molecular Biology Organization; MSU: Michigan State University \*at the time of the award; publication age defined as (2025-first publication year +1), award publication age is (award year-first publication year +1)

86.9% of the 773 Stanford endowed professors, 83.7% of the 129 MSU endowed professors, and 93.8% of the 112 MSU distinguished professors passed the productivity criteria and there was large variability in this regard depending on the school/college involved, e.g. ranging from 100% for Stanford sustainability and education schools and 243/245 (99.2%) for Stanford Medicine (2/245 were renowned educators), to 76.3% for Stanford humanities and sciences and 57.1% for MSU business, again reflecting different types of merit being recognized.

Turing (n=25) and Nobel (n=166) laureates always met the productivity criteria for further analysis with three exceptions: Robert Metcalfe, Peter Higgs, and Akira Yoshino, whose Scopus records included 17, 10, and 19 items, respectively, below the 20-item threshold.

Among the members passing the productivity criteria and thus eligible for analysis (Table 1), the median publication ages for the groups of academies and honorific societies ranged from 22 to 39, with the lowest median ages for ASCI (22 years) and EMBO (25 years), two honorific groups that are known to elect relatively younger scientists, and the highest median ages for NAE (39 years) and NAS (38 years). Except for ASCI and EMBO, the 25^th^ percentile required at least 23 years of published work for election. Cohorts of Stanford and MSU endowed professors had similar median publication ages (28 to 40 years) to those of cohorts of academies and honorific societies. We did not have data on how many years these endowed professors already held their professorships. Turing and Nobel laureates had the highest publication age (median 48 to 55 years). Using each laureate’s first Scopus-indexed publication year and award year, median publication age at the award year was 46 years for Chemistry, 40 for Economics, 44 for Medicine, and 48 for Physics Nobel laureates, and 43 years for Turing laureates.

### Percentile placement for different metrics

As shown in Table 2, the eligible members of honorific groups had modestly good median percentile values for total number of published items, but they were not exceptionally outstanding: the median was never below the 0.5% percentile for any of the 32 groups except for Chemistry Nobel laureates, while endowed professors in business or law had median values in the 9-12% percentile and EMBO members in the 5% percentile. Median percentile values for total citations were good, but not exceptionally outstanding, with medians below 0.5% for NAE, NAS, Stanford engineering and sustainability, and the Turing and Nobel laureates. Total publications were never among the two best-performing metrics in any group. Total citations were best for the Economics Nobel laureates and second best in 2 of the 32 groups.

**Table 2.**
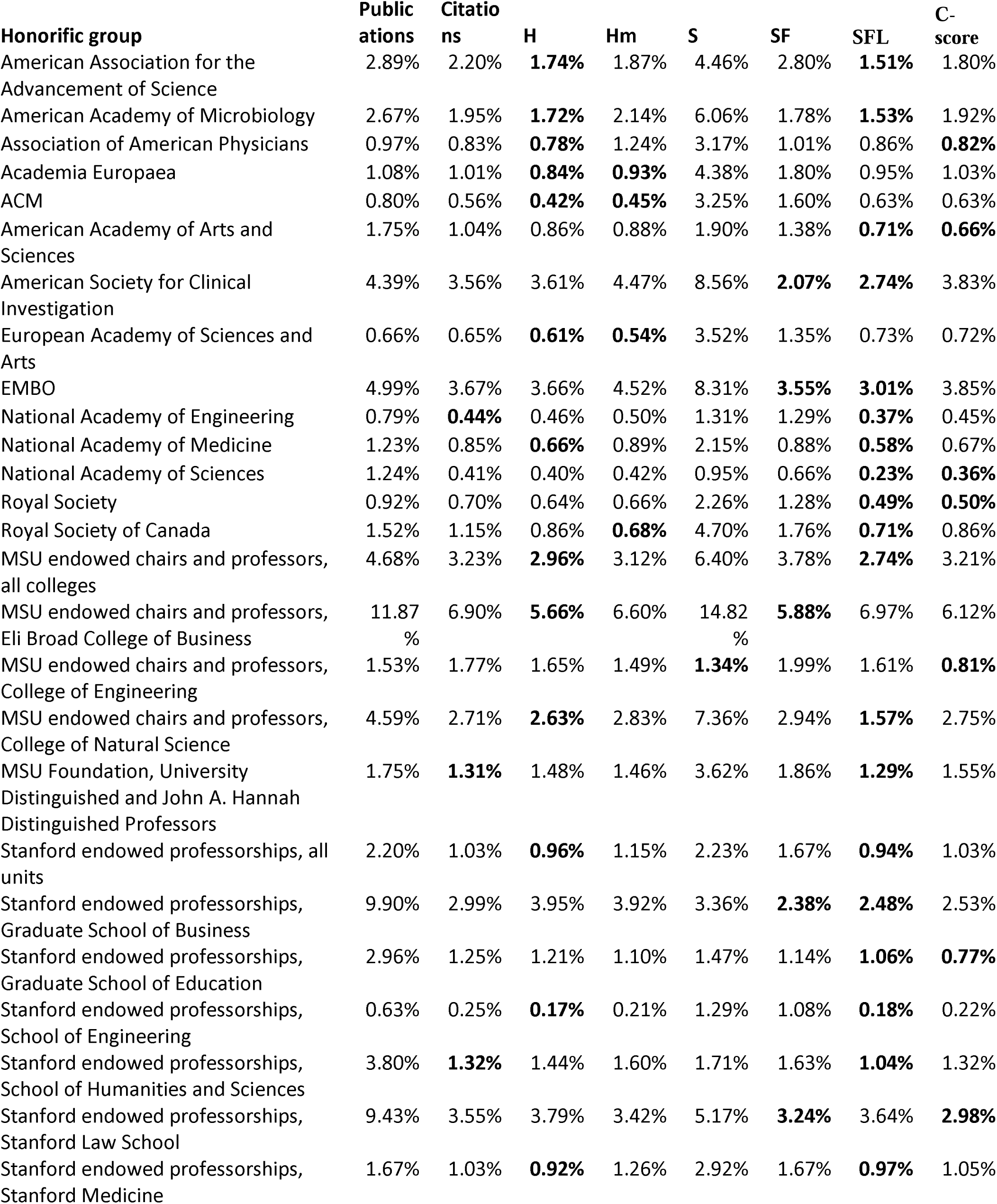
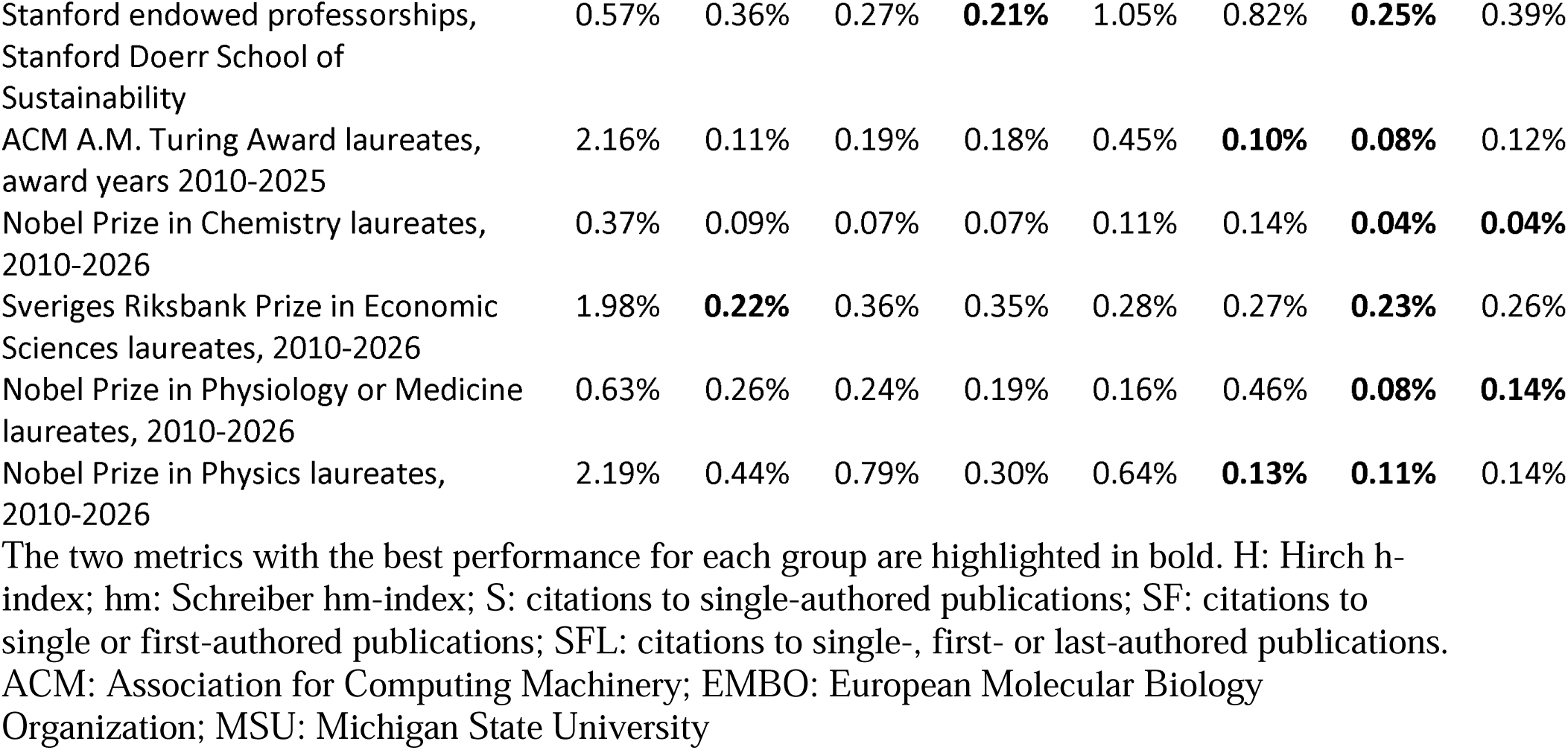
Median percentile (in subfield) for metrics among eligible honorees.

The most frequent metric to have the best performance (lowest median percentile) was citations to single-, first-, and last-authored publications (SFL). It was best for 16 groups (AAAS, AAM, EMBO, NAE, NAM, NAS, Royal Society, MSU all colleges, MSU Natural Science, MSU distinguished professors, Stanford all units, Stanford Humanities and Sciences, Turing, and Nobel laureates in Chemistry, Medicine, and Physics) and second best in another 9 groups. The Hirsch h-index was best for 6 groups (AAP, Academia Europaea, ACM, MSU Business, Stanford Engineering, and Stanford Medicine) and second best for 7. The composite C-score was best for 4 groups (AmAAS, MSU Engineering, Stanford Education, and Stanford Law) and second best for 6. The co-authorship-adjusted Schreiber hm-index was best for 3 groups (EASA, RSC, and Stanford Sustainability) and second best for 2. Citations to single-and first-authored papers were best for 2 groups (ASCI and Stanford Business) and second best for 5. Total citations were best for the Economics Nobel laureates and second best for 2 groups. Citations to single-authored papers were not best for any group and second best for 1.

Across all 32 groups, the best median of the median values was for the c-score (0.81% percentile), followed by the h-index (0.86%), and the number of citations to single-, first-, and last-authored papers (0.90%).

### Inclusion among the top-cited scientists of the Stanford/Elsevier listing

As shown in Table 3, the percentage of eligible members of the honorific groups who appeared among the top-cited scientists of the Stanford/Elsevier listing varied substantially across groups and followed the anticipated gradient of relative prestige. All Turing laureates and all but two Nobel laureates (Mary E. Brunkow (Nobel Prize in Medicine) and Andrea Ghez (Nobel Prize in Physics)) were listed, and coverage in the Stanford/Elsevier listing exceeded 90% also for NAS, AAP and Stanford engineering professorships. The most prestigious official national academies (the three national US academies and the UK Royal Society) had 84-92% coverage, while other honorific groups had 51-76% coverage, except for AAP and ACM that had very high coverage. Honorific societies electing relatively younger members (ASCI, EMBO) had low coverage in the career-wide listing, but better coverage in the single recent year impact listing. Among endowed professorships, MSU distinguished professorships had higher coverage (72%) than MSU professorships (52%), and Stanford professorships had higher coverage (78%) than MSU professorships. Stanford also had better coverage than MSU within comparable school/colleges disciplines. The lowest coverage was seen for business and law disciplines (lowest, 17% for MSU business).

**Table 3.**
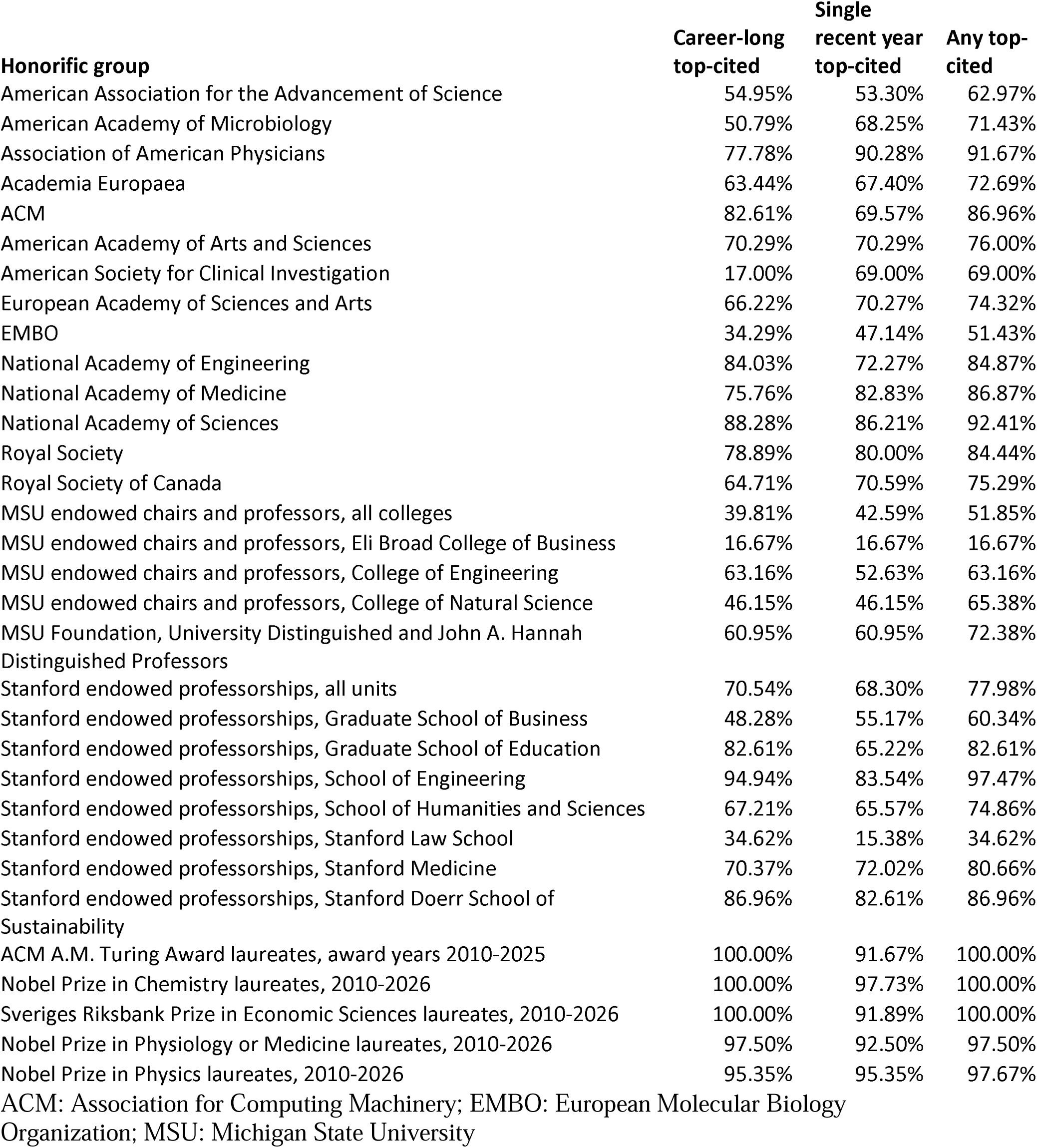
Eligible honorees included in Stanford/Elsevier lists of top-cited authors.

| **Honorific group** | **Career-long top-cited** | **Single recent year top-cited** | **Any top-cited** |
|---|---|---|---|
| American Association for the Advancement of Science | 54.95% | 53.30% | 62.97% |
| American Academy of Microbiology | 50.79% | 68.25% | 71.43% |
| Association of American Physicians | 77.78% | 90.28% | 91.67% |
| Academia Europaea | 63.44% | 67.40% | 72.69% |
| ACM | 82.61% | 69.57% | 86.96% |
| American Academy of Arts and Sciences | 70.29% | 70.29% | 76.00% |
| American Society for Clinical Investigation | 17.00% | 69.00% | 69.00% |
| European Academy of Sciences and Arts | 66.22% | 70.27% | 74.32% |
| EMBO | 34.29% | 47.14% | 51.43% |
| National Academy of Engineering | 84.03% | 72.27% | 84.87% |
| National Academy of Medicine | 75.76% | 82.83% | 86.87% |
| National Academy of Sciences | 88.28% | 86.21% | 92.41% |
| Royal Society | 78.89% | 80.00% | 84.44% |
| Royal Society of Canada | 64.71% | 70.59% | 75.29% |
| MSU endowed chairs and professors, all colleges | 39.81% | 42.59% | 51.85% |
| MSU endowed chairs and professors, Eli Broad College of Business | 16.67% | 16.67% | 16.67% |
| MSU endowed chairs and professors, College of Engineering | 63.16% | 52.63% | 63.16% |
| MSU endowed chairs and professors, College of Natural Science | 46.15% | 46.15% | 65.38% |
| MSU Foundation, University Distinguished and John A. Hannah Distinguished Professors | 60.95% | 60.95% | 72.38% |
| Stanford endowed professorships, all units | 70.54% | 68.30% | 77.98% |
| Stanford endowed professorships, Graduate School of Business | 48.28% | 55.17% | 60.34% |
| Stanford endowed professorships, Graduate School of Education | 82.61% | 65.22% | 82.61% |
| Stanford endowed professorships, School of Engineering | 94.94% | 83.54% | 97.47% |
| Stanford endowed professorships, School of Humanities and Sciences | 67.21% | 65.57% | 74.86% |
| Stanford endowed professorships, Stanford Law School | 34.62% | 15.38% | 34.62% |
| Stanford endowed professorships, Stanford Medicine | 70.37% | 72.02% | 80.66% |
| Stanford endowed professorships, Stanford Doerr School of Sustainability | 86.96% | 82.61% | 86.96% |
| ACM A.M. Turing Award laureates, award years 2010-2025 | 100.00% | 91.67% | 100.00% |
| Nobel Prize in Chemistry laureates, 2010-2026 | 100.00% | 97.73% | 100.00% |
| Sveriges Riksbank Prize in Economic Sciences laureates, 2010-2026 | 100.00% | 91.89% | 100.00% |
| Nobel Prize in Physiology or Medicine laureates, 2010-2026 | 97.50% | 92.50% | 97.50% |
| Nobel Prize in Physics laureates, 2010-2026 | 95.35% | 95.35% | 97.67% |

ACM: Association for Computing Machinery; EMBO: European Molecular Biology Organization; MSU: Michigan State University

Table 4 shows the coverage in the Stanford/Elsevier listing for members of each honorific group according to publication age. Coverage was modest for those with fewer than 20 years of published work, while it was high for those with 20-40 years of published work, and very high for those with more than 40 years of published work. Gradients between the most prestigious and other academies/societies weakened after the publication age stratification, suggesting that part of the difference was due to election at different career stages.

**Table 4.** Inclusion in the Stanford/Elsevier lists of top-cited authors, by publication age.

| **Honorific group** | **<20 years,<br>n/N (%)** | **20-40 years,<br>n/N (%)** | **>40 years,<br>n/N (%)** |
|---|---|---|---|
| American Association for the Advancement of Science | 9/18 (50.0%) | 212/328 (64.6%) | 46/78 (59.0%) |
| American Academy of Microbiology | 2/4 (50.0%) | 38/54 (70.4%) | 5/5 (100.0%) |
| Association of American Physicians | 3/4 (75.0%) | 53/58 (91.4%) | 10/10 (100.0%) |
| Academia Europaea | 9/22 (40.9%) | 118/162 (72.8%) | 38/43 (88.4%) |
| ACM | 2/3 (66.7%) | 55/63 (87.3%) | 3/3 (100.0%) |
| American Academy of Arts and Sciences | 4/11 (36.4%) | 79/106 (74.5%) | 50/58 (86.2%) |
| American Society for Clinical Investigation | 24/34 (70.6%) | 44/65 (67.7%) | 1/1 (100.0%) |
| European Academy of Sciences and Arts | 3/10 (30.0%) | 45/56 (80.4%) | 7/8 (87.5%) |
| EMBO | 8/16 (50.0%) | 27/53 (50.9%) | 1/1 (100.0%) |
| National Academy of Engineering | 2/2 (100.0%) | 63/75 (84.0%) | 36/42 (85.7%) |
| National Academy of Medicine | 4/8 (50.0%) | 67/76 (88.2%) | 15/15 (100.0%) |
| National Academy of Sciences | 3/3 (100.0%) | 79/86 (91.9%) | 52/56 (92.9%) |
| Royal Society | 0/0 (n/a) | 57/70 (81.4%) | 19/20 (95.0%) |
| Royal Society of Canada | 1/3 (33.3%) | 54/71 (76.1%) | 9/11 (81.8%) |
| MSU endowed chairs and professors, all colleges | 7/22 (31.8%) | 37/71 (52.1%) | 12/15 (80.0%) |
| MSU endowed chairs and professors, Eli Broad College of Business | 0/2 (0.0%) | 1/9 (11.1%) | 1/1 (100.0%) |
| MSU endowed chairs and professors, College of Engineering | 1/4 (25.0%) | 10/13 (76.9%) | 1/2 (50.0%) |
| MSU endowed chairs and professors, College of Natural Science | 4/7 (57.1%) | 8/14 (57.1%) | 5/5 (100.0%) |
| MSU Foundation, University Distinguished and John A. Hannah Distinguished Professors | 8/10 (80.0%) | 45/62 (72.6%) | 23/33 (69.7%) |
| Stanford endowed professorships, all units | 22/42 (52.4%) | 326/433 (75.3%) | 176/197 (89.3%) |
| Stanford endowed professorships, Graduate School of Business | 3/8 (37.5%) | 25/41 (61.0%) | 7/9 (77.8%) |
| Stanford endowed professorships, Graduate School of Education | 1/3 (33.3%) | 13/15 (86.7%) | 5/5 (100.0%) |
| Stanford endowed professorships, School of Engineering | 0/0 (n/a) | 54/54 (100.0%) | 23/25 (92.0%) |
| Stanford endowed professorships, School of Humanities and Sciences | 6/8 (75.0%) | 74/109 (67.9%) | 57/66 (86.4%) |
| Stanford endowed professorships, Stanford Law School | 0/2 (0.0%) | 8/21 (38.1%) | 1/3 (33.3%) |
| Stanford endowed professorships, Stanford Medicine | 11/19 (57.9%) | 122/157 (77.7%) | 63/67 (94.0%) |
| Stanford endowed professorships, Stanford Doerr School of Sustainability | 0/1 (0.0%) | 10/11 (90.9%) | 10/11 (90.9%) |
| ACM A.M. Turing Award laureates, award years 2010-2025 | 0/0 (n/a) | 2/2 (100.0%) | 22/22 (100.0%) |
| Nobel Prize in Chemistry laureates, 2010-2026 | 2/2 (100.0%) | 7/7 (100.0%) | 35/35 (100.0%) |
| Sveriges Riksbank Prize in Economic Sciences laureates, 2010-2026 | 0/0 (n/a) | 8/8 (100.0%) | 29/29 (100.0%) |
| Nobel Prize in Physiology or Medicine laureates, 2010-2026 | 0/0 (n/a) | 7/8 (87.5%) | 32/32 (100.0%) |
| Nobel Prize in Physics laureates, 2010-2026 | 0/0 (n/a) | 7/8 (87.5%) | 35/35 (100.0%) |

ACM: Association for Computing Machinery; EMBO: European Molecular Biology Organization; MSU: Michigan State University

### Research integrity proxies

As shown in Table 5, all academies and honorific societies had at least 1 or more members in 2026 who had a retraction history, with rates ranging from 1.1% (Royal Society) to 9.5% (AAM). MSU professorships included only one person with a retraction history (0.9%) and MSU distinguished professors included no one with retractions, while Stanford endowed professors had a substantial percentage (4.9%) with retraction history. Turing and Economics Nobel laureates did not have any retractions, and only one Physics Nobel laureate had a retraction history. Retraction history was common in Chemistry Nobel laureates (11.4%) and even more in Medicine Nobel laureates (17.5%). Honored individuals with high self-citations (above the 95^th^ percentile among top-cited scientists in the same subfield) were extremely uncommon (only 2 such individuals were identified). A few honored individuals had high proportion of publications in Scopus-discontinued titles, especially in NAE (3.4%), AE (2.6%), Stanford engineering (2.5%), AmAAS (1.7%), and Stanford Medicine (1.7%); and this was also observed in a Physics Nobel laureate, Klaus L. Hasselmann.

**Table 5.** Research integrity proxies.

| Honorific group | Eligible members | Retractions (%) | High self-citations (%) | High discontinued titles (%) |
|---|---|---|---|---|
| American Association for the Advancement of Science | 424 | 15 (3.54%) | 0 (0.00%) | 7 (1.65%) |
| American Academy of Microbiology | 63 | 6 (9.52%) | 1 (1.59%) | 1 (1.59%) |
| Association of American Physicians | 72 | 4 (5.56%) | 0 (0.00%) | 0 (0.00%) |
| Academia Europaea | 227 | 9 (3.96%) | 0 (0.00%) | 6 (2.64%) |
| ACM | 69 | 3 (4.35%) | 0 (0.00%) | 0 (0.00%) |
| American Academy of Arts and Sciences | 175 | 10 (5.71%) | 0 (0.00%) | 3 (1.71%) |
| American Society for Clinical Investigation | 100 | 2 (2.00%) | 1 (1.00%) | 0 (0.00%) |
| European Academy of Sciences and Arts | 74 | 5 (6.76%) | 0 (0.00%) | 1 (1.35%) |
| EMBO | 70 | 4 (5.71%) | 0 (0.00%) | 1 (1.43%) |
| National Academy of Engineering | 119 | 2 (1.68%) | 0 (0.00%) | 4 (3.36%) |
| National Academy of Medicine | 99 | 4 (4.04%) | 0 (0.00%) | 1 (1.01%) |
| National Academy of Sciences | 145 | 8 (5.52%) | 0 (0.00%) | 2 (1.38%) |
| Royal Society | 90 | 1 (1.11%) | 0 (0.00%) | 1 (1.11%) |
| Royal Society of Canada | 85 | 1 (1.18%) | 0 (0.00%) | 0 (0.00%) |
| MSU endowed chairs and professors, all colleges | 108 | 1 (0.93%) | 0 (0.00%) | 0 (0.00%) |
| MSU endowed chairs and professors, Eli Broad College of Business | 12 | 0 (0.00%) | 0 (0.00%) | 0 (0.00%) |
| MSU endowed chairs and professors, College of Engineering | 19 | 0 (0.00%) | 0 (0.00%) | 0 (0.00%) |
| MSU endowed chairs and professors, College of Natural Science | 26 | 0 (0.00%) | 0 (0.00%) | 0 (0.00%) |
| MSU Foundation, University Distinguished and John A. Hannah Distinguished Professors | 105 | 0 (0.00%) | 0 (0.00%) | 1 (0.95%) |
| Stanford endowed professorships, all units | 672 | 33 (4.91%) | 0 (0.00%) | 8 (1.19%) |
| Stanford endowed professorships, Graduate School of Business | 58 | 1 (1.72%) | 0 (0.00%) | 0 (0.00%) |
| Stanford endowed professorships, Graduate School of Education | 23 | 0 (0.00%) | 0 (0.00%) | 0 (0.00%) |
| Stanford endowed professorships, School of Engineering | 79 | 3 (3.80%) | 0 (0.00%) | 2 (2.53%) |
| Stanford endowed professorships, School of Humanities and Sciences | 183 | 6 (3.28%) | 0 (0.00%) | 2 (1.09%) |
| Stanford endowed professorships, Stanford Law School | 26 | 0 (0.00%) | 0 (0.00%) | 0 (0.00%) |
| Stanford endowed professorships, Stanford Medicine | 243 | 20 (8.23%) | 0 (0.00%) | 4 (1.65%) |
| Stanford endowed professorships, Stanford Doerr School of Sustainability | 23 | 1 (4.35%) | 0 (0.00%) | 0 (0.00%) |
| ACM A.M. Turing Award laureates, award years 2010-2025 | 24 | 0 (0.00%) | 0 (0.00%) | 0 (0.00%) |
| Nobel Prize in Chemistry laureates, 2010-2026 | 44 | 5 (11.36%) | 0 (0.00%) | 0 (0.00%) |
| Sveriges Riksbank Prize in Economic Sciences laureates, 2010-2026 | 37 | 0 (0.00%) | 0 (0.00%) | 0 (0.00%) |
| Nobel Prize in Physiology or Medicine laureates, 2010-2026 | 40 | 7 (17.50%) | 0 (0.00%) | 0 (0.00%) |
| Nobel Prize in Physics laureates, 2010-2026 | 43 | 1 (2.33%) | 0 (0.00%) | 1 (2.33%) |

ACM: Association for Computing Machinery; EMBO: European Molecular Biology Organization; MSU: Michigan State University; high self-citations and high discontinued titles are defined as having a proportion of self-citations or proportion of publications in discontinued titles above the 95<sup>th</sup> percentile, among top-cited author profiles in their subfield.

## DISCUSSION

We analyzed data on 32 groups of scientists receiving major and extraordinary recognitions. These are all groups including mostly scholars with a long track record, but they vary considerably in the average publication age of the honorees, ranging from relatively younger scientists (e.g. ASCI) to mostly very senior ones (e.g. Turing and Nobel laureates). We found that total number of published items and total citations did not have that good performance in identifying the average member of these honorific groups. Among the tested metrics, the number of citations to single-, first-or last-authored publications had the best performance for the average honoree in 16/32 groups based on the median percentile value, while the composite c-score had the best median of median percentile values across the 32 groups. Some other metrics also did well in specific groups. The percentage of members who were covered by the Stanford/Elsevier listing of top-cited scientists was congruent with the relative perceived prestige of these recognitions; more prestigious recognitions had higher coverage, reaching as high as 99% for Turing and Nobel laureates. Research integrity proxies showed that a substantial number of these honorees have a retraction history, with highest occurrences for Medicine and Chemistry Nobel laureates; a few have published many publications in Scopus-discontinued titles; and high percentages of self-citations were almost never observed.

One may argue as to what extent the bestowing of these coveted recognitions represents a platinum, gold or silver standard of merit or excellence. Moreover, should metrics benchmark recognitions or recognitions benchmark metrics? Scientific and scholarly impact can be extremely multifarious, and these recognitions may reward predominantly some phenotypes rather than others. Even the awards that supposedly cover vast scientific areas in the broadest terms may, in reality, get concentrated in relatively few disciplines (23) and types of contributions. Similarly, scientometrics only capture specific dimensions of impact. Given this unavoidable restriction in the face validity of both recognitions and citation metrics, it is even more remarkable how closely some of these metrics track the average honored scholar and even provide a gradient that is highly congruent with the perceived relative prestige of these recognitions. This may be interpreted in various ways: metrics may genuinely have strong association with and/or be strong markers of scholarly merit; or, conversely, selection for these honors may be heavily guided by metrics: evaluators may be using metrics heavily for deciding the award winners, inductees, and endowed chairs. Regardless, our work cannot assess why some other individuals with similar or perhaps even better metrics do not receive these honors. Names of nominees for major awards and for academies and honorific societies are not promptly available in public view, and it is difficult to conceive a suitable control group of people who are considered for but not granted endowed professorships.

Regardless of the challenges of etiological attribution, some metrics seem to be better markers than others. Citations to single-, first-or last-authored publication was the metric with the best median percentile values for half of the examined groups. In most disciplines, these authorship positions signify the lion’s share in the conception, execution, and presentation of the work (24–26). Not surprisingly, this metric is dominant in the most prestigious awards, where one seeks to select the individuals with the largest contribution to major discoveries and achievements. However, author position seems to be pervasively important across honorific recognitions and even in younger cohorts. In ASCI, in particular, which is the youngest of the honorific societies examined here, citations to single-or first-authored publications had the best median percentile values. Conversely, total number of publications and total number of citations had relatively unimpressive performance in these honorific groups. This is welcome, since number of publications has become a highly gameable indicator (27–29) and even total number of citations may be gamed to some extent (30). However, there is a need to recognize also diverse contributions which may not result in leading authorship positions but are indispensable for good science, especially team science (31), to materialize. For many top recognitions, team science has been difficult or even impossible to recognize. Changing the reward systems to give more value to collaborative work nevertheless needs to be balanced against the higher gaming potential of popular, but also widely misused metrics such as the h-index that give full credit to all members of large teams but ignore relative contributions in each paper. The wide use of such gameable metrics may promote unwarranted inflation of authorship and gift authorship (32,33). Appropriate collective credit allocation in multi-authorship (34) that would be feasible at science-wide scale remains highly desirable but very difficult to achieve with fairness and transparency.

The composite c-score had the best median of median percentiles across the 32 groups. It provides a metric that behaves predictably well across very diverse important, major, and extreme academic recognitions. It may be more difficult than raw citations or h-index, but not impossible (30), to reach the top in this composite metric through gaming or fraud. More importantly, the Stanford/Elsevier listing of top-cited scientists captured different proportions of honored individuals in proportionality to the magnitude of the bestowed recognition. Of course, individual variability can be large, and exceptions may occur even for the most prestigious honors. For example, two Nobel laureates were not included in either the career-long or single-year Stanford/Elsevier listings for 2026 and it is possible that additional ones might not qualify for these listings at the time of the Nobel prize announcement. Previous work has shown diversity in career trajectories in Nobel laureates and their impact (35). However, hot streaks with high citations (35), and a boosting of citations after the award (36) are very common hallmarks. Even if there is a dip in subsequent productivity after the award (37), very high cumulative impact is a largely ubiquitous feature for laureates. In general, recognitions given to scientists in their early and mid-career may precede the accumulation of major citation impact; while accumulated major citation impact is more certain for those in more senior phases of their career.

For these cohorts of influential scholars, it is not surprising that self-citations remain at reasonable, modest levels. Moreover, high proportion of publications in titles that were subsequently discontinued from Scopus indexing were not a common feature. Even when observed, this may not necessarily have reflected the reputation of these journals when the honorees published their work in them. Discontinuation may have happened much later when the journal’s reputation had changed. For example, four papers by Nobel laureate Klaus Hasselmann were published in Physics Essays 1996–1997, whereas Scopus coverage continued through 2017. That journal was launched in 1988 with the laudable goal to publish peer-reviewed work on alternative mathematical frameworks that challenged the mainstream. However, Scopus and other indexing databases stopped covering it ∼30 years later; by then it had become occupied too much with what many might perceive as pseudoscience.

Retractions were the most common research integrity proxy indicator that we observed across the analyzed honorees. They were most prominent in Medicine and Chemistry Nobel laureates. They may reflect the extra scrutiny that the work of these individuals receives after they reach that pinnacle of honor. Interestingly, all academies and honorific societies inducted in 2026 some member(s) or fellow(s) with a history of retractions and Stanford professors had more frequent history of retractions than MSU professors. Thus, retractions may follow the same gradient as the perceived prestige gradient. Retractions may not necessarily reflect misconduct, but honest error (38). When authors retract their work after they recognize error, retractions should be applauded. However, the exact cause is often not clearly reported in retraction notices (39) and most surveys suggest that misconduct is the most common reason for retractions (40,41). More attention to research integrity proxies may be needed to achieve a better balance of impact and integrity in appraising merit.

Some limitations of our work should be noted. First, Scopus has high precision and recall (12,42), but some errors in author files do exist and they may affect the profiles and percentile rankings of single individuals; and we cannot exclude some errors in matching individuals to a specific Scopus author ID file. Nevertheless, we carefully reviewed results and made some additional manual corrections, so residual errors should be small. Second, we used an arbitrary productivity threshold to exclude individuals who were likely honored for reasons not predominantly related to their published scientific work and its impact. Occasionally some scientists may publish one or very few, but extremely influential papers, and little else in their career. However, this phenotype represented by the legendary examples of Robert Metcalfe (Ethernet discovery), Peter Higgs (Higgs boson), and Akira Yoshino (practical lithium-ion batteries) has become rare. Moreover, there can be no consensus on what proportion of academic honors should go to individuals for contributions other than what is reflected in scholarly published work. As we have mapped in this analysis, different honorific organizations have very different traditions in this regard. Fourth, for endowed professors and Turing and Nobel laureates we captured their current metrics rather than those they had when they were given these recognitions, while for academies and honorific societies announced in 2026, citation metrics through 2025 generally preceded the time of induction. Fifth, we gave precedence to the median values of the percentile metrics, to focus on the average member of each of these prestigious groups. Nevertheless, the range of values can be very large across the members of each group. Therefore, under no circumstances should our results be interpreted that the scientists above the median values deserve some recognition and those below do not deserve it or necessarily deserve it less. Moreover, subtle differences in these metrics are unlikely to be informative.

We speculate that metrics are probably already influencing to various extent the selection of individuals for these recognitions. The appropriateness or inappropriateness of this use is hard to decipher without knowing the full deliberations of the honoring body in each scientist’s case. Regardless, the average values of these metrics and the proportion of top-cited scientists track remarkably well the perceived prestige of the examined recognitions.

## Data

Full data is available in the August 2026 iteration of https://elsevier.digitalcommonsdata.com/datasets/btchxktzyw/9 (version 9) and in the Supplementary Tables 1 and 2

